# *URA6* mutations provide an alternative mechanism for 5-FOA resistance in *Saccharomyces cerevisiae*

**DOI:** 10.1101/2024.06.03.597250

**Authors:** Joseph O. Armstrong, Pengyao Jiang, Skyler Tsai, Megan My-Ngan Phan, Amy A. Caudy, Kelley Harris, Maitreya J. Dunham

**Author notes:** Contributed equally (Joseph O. Armstrong, Pengyao Jiang).

## Abstract

The *URA3* gene is frequently used in the budding yeast community as the mutation target for 5-fluoroorotic acid (5-FOA) resistance. We identified a class of *ura6* mutants that can grow in the presence of 5-FOA. Unlike *ura3* mutants, *ura6* mutants remain prototrophic and are able to grow in the absence of uracil. In addition to 5-FOA resistance, we found that *ura6* mutants are also resistant to 5-fluorocytosine (5-FC) and 5-fluorouracil (5-FU). In total, we identified 50 unique missense mutations across 32 unique amino acid positions of Ura6 which confer resistance to 5-FOA. We found that 28 out of the 32 affected positions are located in regions conserved between *Saccharomyces cerevisiae* and three clinically relevant pathogenic fungi. Metabolic analysis revealed a build-up of uridine monophosphate (UMP) and 5-fluorouridine monophosphate (5-FUMP) in *ura6* mutants, indicating a reduction in Ura6 activity. Despite the accumulation of UMP in *ura6* mutants, uridine diphosphate and triphosphate (UDP, UTP) levels were similar across mutants and wild type. These findings suggest that missense mutations to *URA6*, can result in reduction of uridylate kinase activity and lead to cross resistance to fluorinated prodrugs that incorporate into both the *de novo* pyrimidine synthesis and the pyrimidine salvage pathway.

## Introduction

The *URA3* locus is a versatile tool for yeast geneticists and is used in a wide range of applications involving plasmid transformation, recombination, and mutation assays in *Saccharomyces cerevisiae* and other fungi (Lang and Murray 2008; Kaneko et al. 2009; Boeke, LaCroute, and Fink 1984). *URA3* is particularly useful because it can be positively selected for by growing cells in the absence of uracil, or selected against by growing cells in the presence of 5-fluoroorotic acid (5-FOA). Orotic acid (OA) is a metabolite in the *de novo* pyrimidine synthesis pathway upstream of Ura3. Both OA and 5-FOA are metabolized by the major and minor forms of orotate phosphoribosyltransferases encoded by *URA5* and *URA10* to form OMP and 5-fluoroorotidine monophosphate (5-FOMP), respectively. 5-FOMP is metabolized into 5-fluorouridine monophosphate (5-FUMP). The toxicity of 5-FOA is due to the subsequent conversion of 5-FUMP to 5-fluorouridine triphosphate (5-FUTP) (Longley, Harkin, and Johnston 2003; Vermes, Guchelaar, and Dankert 2000). Cells with a loss-of-function mutation to *URA3* are resistant to 5-FOA because 5-FOMP can no longer be metabolized into the toxic downstream metabolites (Boeke, LaCroute, and Fink 1984). A loss-of-function mutation to *URA3* ablates the *de novo* pyrimidine synthesis pathway, and *ura3* mutants must therefore salvage uracil from the environment to grow.

*URA3* has been used as a mutation reporter in yeast genetics research for several decades (Lang and Murray 2008, 2011; Kiktev et al. 2018; Williams et al. 2021; Zhou et al. 2021; Lee et al. 1988). Although *URA3* is well known in the literature to be the primary target for 5-FOA resistance, there are unexplained instances where cells containing a wild-type copy of *URA3* were resistant to 5-FOA. For example, 10% of mutants that were discovered through a fluctuation assay contained a wild-type copy of *URA3* (Lang and Murray 2008). More astonishingly, Boeke et al. reported that only 5-10% of a set of UV-induced 5-FOA mutants contained mutations to *URA3* (Boeke, LaCroute, and Fink 1984). The genetic cause of these 5-FOA-resistant cells containing a wild-type copy of *URA3* remains unknown.

When we sought to use *URA3* as a mutation reporter in *S. cerevisiae*, we unexpectedly found a large number of mutant colonies containing wild-type *URA3* that readily grew in the presence of 5-FOA. These mutants also grew readily in the absence of uracil, confirming that the pyrimidine *de novo* synthesis pathway remained functional. Using whole genome sequencing, we identified that all resistant colonies with wild-type *URA3* had mutations to *URA6*, which has been previously linked to 5-FOA resistance, but not thoroughly characterized (Nivedita, Aitchison, and Baliga 2021; McAtee 2013). In addition to 5-FOA resistance, we also found that *ura6* mutants are resistant to fluorinated pyrimidine derivatives such as 5-fluorocytosine (5-FC) and 5-fluorouracil (5-FU). We analyzed metabolites from wild-type and *ura6* mutants grown in both minimal media and minimal media containing 5-FOA and found that UMP and 5-FUMP levels are increased in *ura6* mutants, indicating that a reduction of function of Ura6 drives resistance to fluorinated pyrimidine derivatives. It is not immediately apparent how *ura6* mutants retain sufficient pyrimidine biosynthesis to support growth while simultaneously avoiding the toxicity of fluorinated pyrimidine analogs. We compared the metabolome of WT and *ura6* mutants and found that *ura6* mutants build up UMP indicating that these mutations lead to a reduction of flux through Ura6. We propose a model for the mechanism of resistance where the build-up of UMP competitively inhibits the metabolism of 5-FUMP which in turn leads to resistance. To our knowledge, *URA6* is the sole mutable target capable of providing cross resistance to fluorinated prodrugs incorporated into both the pyrimidine synthesis and salvage pathways. Mutations to *URA6* represent a novel mechanism of cross-resistance to fluorinated pyrimidine analogs. Our findings parallel a recent report describing a similar class of *ura6* mutants in *Schizosaccharomyces pombe* (Kowal et al. 2026).

## Results

### Discovery of a class of ura6 mutants which confers 5-FOA resistance

We discovered a new class of mutants that can grow in 5-FOA medium with the wildtype (WT) *URA3* allele in a haploid strain. We plated a large number of cells (on the order of 10^8^ for each sample) onto 5-FOA with the intention of identifying *de novo* mutations to *URA3*. The initial inoculants were grown in synthetic complete media without uracil (SC-ura) media to exclude all mutations that occurred in the *URA3* gene during the overnight growth. We used a pooled sequencing approach aiming to obtain the mutation spectrum of the *URA3* gene, similar to a previously published method using *CAN1* as a reporter gene (Jiang et al. 2021; Jiang, Ollodart, and Dunham 2022). To our surprise, less than half of the total number of mutants pooled in different samples contained mutations to *URA3* (See Methods and Supplementary Figure 1 for details). We performed Sanger sequencing on the *URA3* locus from individual mutant clones contained within two pools and found that only 5 out of 24 clones tested showed mutations in *URA3*. This was surprising because all the mutants we isolated grew stably in the 5-FOA media. We then tested whether the 19 mutants with wild type *URA3* retained a functional pyrimidine synthesis pathway by growing the isolated clones on media without uracil. To our surprise, the isolated mutants grew stably in minimal media as well as in minimal media with 5-FOA (Fig. 1A).

**Figure 1.**
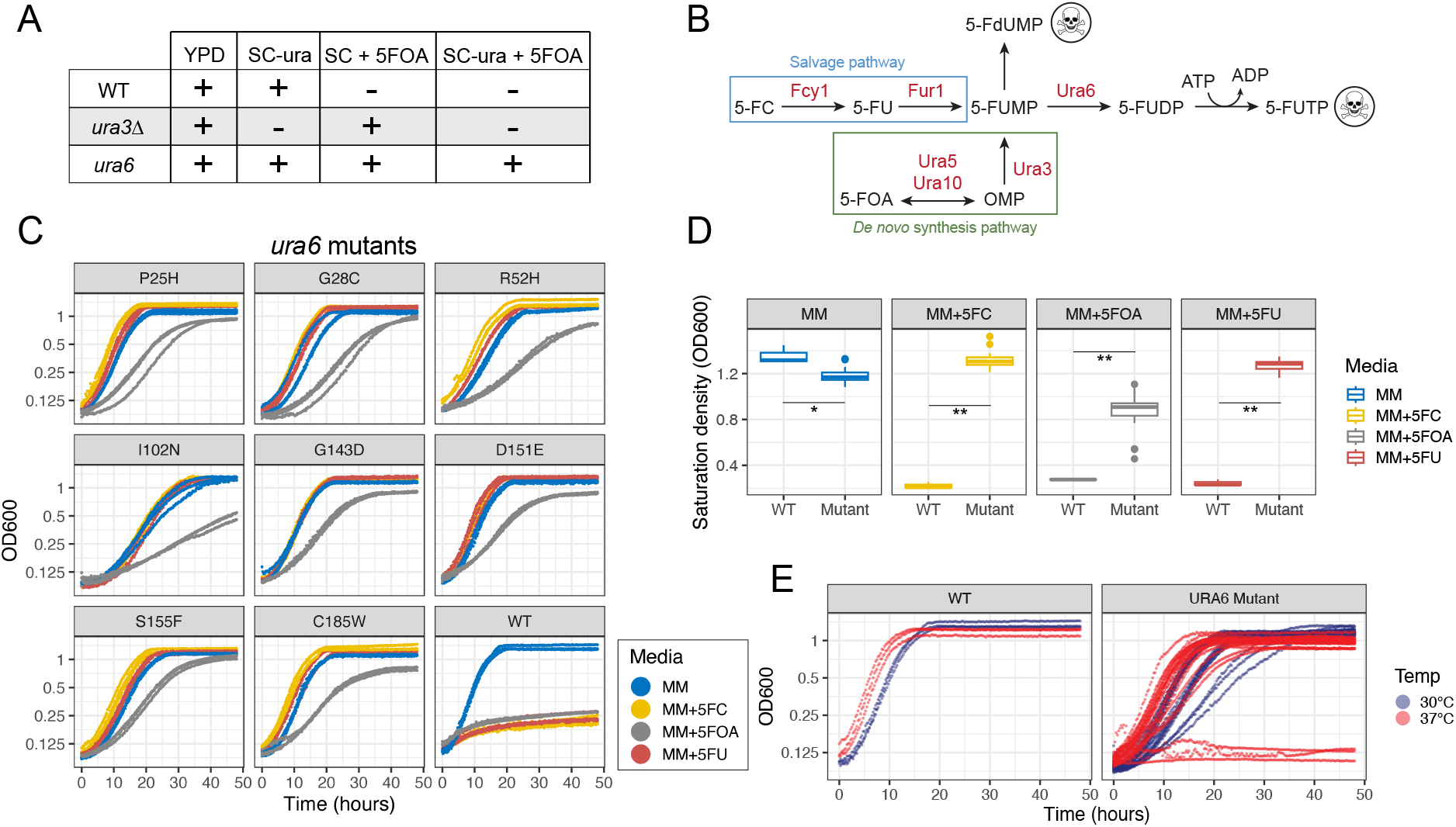
*ura6* mutants are resistant to fluorinated pyrimidine derivatives. A) Comparison of growth phenotypes across WT, *ura3Δ*, and *ura6* mutants in YPD, SC-ura, 5-FOA, SC-ura +5-FOA. “+” indicates growth, and “-” indicates lack of growth (YPD – yeast extract, peptone and dextrose; SC – synthetic complete media). B) The fluorinated drugs 5-FC and 5-FU are incorporated into the pyrimidine salvage pathway. 5-FOA is incorporated into the pyrimidine synthesis pathway. Ura6 lies at the junction of the pyrimidine synthesis and salvage pathways. The production of 5-FUTP from the fluorinated prodrugs leads to toxicity. C) Growth curves for three biological replicates of *ura6* mutants and wild-type in MM (minimal media), MM+5-FOA, MM+5-FC, MM+5-FU measured at 30°C. D) Saturation density for growth for *ura6* mutants and wild-type strains grown in MM, MM+5-FOA, MM+5FC, MM+5FU measured at 30°C. Wilcoxon rank-sum tests with Benjamini-Hochberg correction were used to compare saturation density between wild type and mutant within each media condition. Significance thresholds: *p < 0.05, **p < 0.01. E) Growth curves for three biological replicates of individual *ura6* mutants and wild type strains grown in MM at 30°C and 37°C.

To uncover the genetic basis of the 5-FOA resistant mutants that maintained WT copies of *URA3*, we performed whole genome sequencing of ten individual clones from two independent culture experiments. We found a missense mutation in the coding region of *URA6* in each of the clones we sequenced. These mutations occupy four different sites along the gene (Supplementary Table 1). Five mutants have the same mutation at the 31st amino acid, changing a G to a T, converting a Threonine to an Asparagine. To obtain a more comprehensive distribution of mutations in *URA6* in our initial experiments, we used PCR to amplify the *URA6* locus from previously pooled mutants and performed Illumina sequencing and mutation calling. We recovered an additional 243 *ura6* mutants that gave rise to 5-FOA resistance (Supplementary Figure 2). We identified a total of 50 unique mutations in *URA6*, all of which are missense mutations (Supplementary Table 2). *URA6* is an essential gene, so we predicted that the 50 missense mutations we identified do not result in complete loss of function of the enzyme. We isolated eight individual *ura6* mutants from these pools for further quantitative characterization.

### Mutations to URA6 provide cross resistance to 5-FU and 5-FC in addition to 5-FOA

We initially discovered the *ura6* mutants through their resistance to 5-FOA and their ability to grow in media lacking uracil, a phenotype distinct from *ura3Δ* mutants (Fig. 1A). When incorporated into the cell, fluorinated pyrimidine derivatives converge on a common downstream metabolite, 5-FUMP, which is a substrate of Ura6 and a precursor to the toxic product 5-FUTP. We hypothesized that the *ura6* mutants we identified would also be resistant to 5-FC and 5-FU, since Ura6 operates at the junction of the pyrimidine synthesis and salvage pathways (Fig. 1B). Previous studies found that mutations to *URA6* confer resistance to 5-FC in both pathogenic fungi and *S. cerevisiae* (Durand et al. 2023; Chang et al. 2021). To test whether the *ura6* mutants in our study were resistant to 5-FC and 5-FU in addition to 5-FOA, we employed a plate reader-based growth assay.

We initially discovered the *ura6* mutants through their resistance to 5-FOA and their ability to grow in media lacking uracil, a phenotype distinct from *ura3Δ* mutants (Fig. 1A). When incorporated into the cell, fluorinated pyrimidine derivatives converge on a common downstream metabolite, 5-FUMP, which is a substrate of Ura6 and a precursor to the toxic product 5-FUTP. We hypothesized that the *ura6* mutants we identified would also be resistant to 5-FC and 5-FU, since Ura6 operates at the junction of the pyrimidine synthesis and salvage pathways (Fig. 1B). Previous studies found that mutations to *URA6* confer resistance to 5-FC in both pathogenic fungi and *S. cerevisiae* (Durand et al. 2023; Chang et al. 2021). To test whether the *ura6* mutants in our study were resistant to 5-FC and 5-FU in addition to 5-FOA, we employed a plate reader-based growth assay.

To gain a quantitative understanding of the effects of mutations to *URA6* on growth, we measured growth in the following media: minimal media (MM), MM+5-FOA, MM+5-FU, MM+5-FC (Fig. 1C). In addition, we also measured growth in synthetic complete media (SC), and SC-uracil (Supplementary Figure 3). We found that *ura6* mutants grew similarly in SC and MM which indicated that uracil supplementation had little effect on growth. We used MM as the baseline condition for quantitative comparisons. We compared growth between eight unique *ura6* mutants (n = 3 biological replicates each, 24 total cultures) and the WT strain in each condition (Fig. 1D, Supplementary Fig. 4/5). In minimal media, *ura6* mutants exhibited modestly reduced saturation density relative to WT (mean OD600 : 1.18 vs 1.36, p < 0.05, Fig. 1D). In all three drug conditions (5-FOA, 5-FC, 5-FU) mutants showed a pronounced and consistent growth advantage over wild type (Fig. 1D). Mean saturation density was 1.31 vs 0.22 in MM+5-FC, 1.27 vs 0.25 in MM+5FU, and 0.87 vs 0.28 in MM+5-FOA for *ura6* mutants vs WT.

We calculated maximum specific growth rate (μ_max_) using a rolling 4-hour window of the growth curve (Supplemental Fig. 4). We retained the slope of ln(OD600) with fits R^2^ > 0.98 and μ_max_ was assigned from the median slope of the five steepest windows (Supplemental Fig. 5). Similar to our previous observations of saturation density, *ura6* mutants maintained comparable growth rates in MM, MM+5-FC and MM+5-FU and reduced growth rates in MM + 5-FOA (Supplementary Fig. 5). We observe no tradeoff among *ura6* mutants across fluorinated drug conditions and instead observed a positive correlation between growth in MM and growth in MM + fluorinated prodrug. This contrasts with a recent comprehensive study of *FCY1* mutants in the salvage pathway (Després et al. 2022), which revealed a fitness tradeoff where mutants highly resistant to 5-FC grew poorly in the absence of cytosine.

### ura6 mutants are not exclusively temperature sensitive

Several previously identified *ura6* mutants (e.g. *ura6-1, ura6-4, ura6-5*) were found to be temperature-sensitive (Liljelund and Lacroute 1986; Berg et al. 2020), showing permissive growth at 30°C and thermosensitivity at 37°C. To examine the isolated *ura6* mutants for temperature sensitivity, we measured growth for each mutant in MM and MM+5-FOA. Of the eight individual mutants we isolated for phenotyping, only one (I102N) was unable to grow in MM at 37°C (Fig. 1E). Notably, this I102N mutant had a slower growth rate than other mutants at 30°C in MM, MM+5FC, MM+5FU and MM+5-FOA (Fig. 1C, Supplementary Figure 7). While the I102N mutation confers a temperature sensitivity, we do not observe a difference between the growth of the other isolated mutants at 30°C and 37°C which distinguishes them from previously identified temperature-sensitive alleles (Fig. 1E).

### Mutations to URA6 that cause cross resistance to 5-FC, 5-FU, and 5-FOA typically occur in conserved regions

Evolutionarily conserved sites are commonly used as indicators of functional importance, as strong purifying selection tends to preserve positions that are critical for protein structure or biological function (Cagiada et al. 2023; Capra and Singh 2007; Chakrabarti and Lanczycki 2007). To determine whether the identified mutations are consistent with hypomorphs, we asked whether they occur at evolutionarily conserved positions in Ura6.

In total, we identified 50 unique missense mutations across 32 residues of *URA6* (Fig. 2). We compared the positions of the mutations to domains annotated on UniProt (UniProt Consortium 2023). We found mutations at one of three sites reported to contribute to ATP binding, and four of the six reported ribonucleoside 5’-phosphate binding sites; however, many of the mutations we identified do not overlap with known functional domains. Notably, the distribution of these 50 mutations is not random along the gene (Chi-square test, *p* <0.05, Supplementary Figure 8, robust to different bin sizes). The first ATP binding site and its adjacent sequences, close to amino acid position 25, appear to be especially enriched for mutations that confer resistance to 5-FOA.

**Figure 2.**
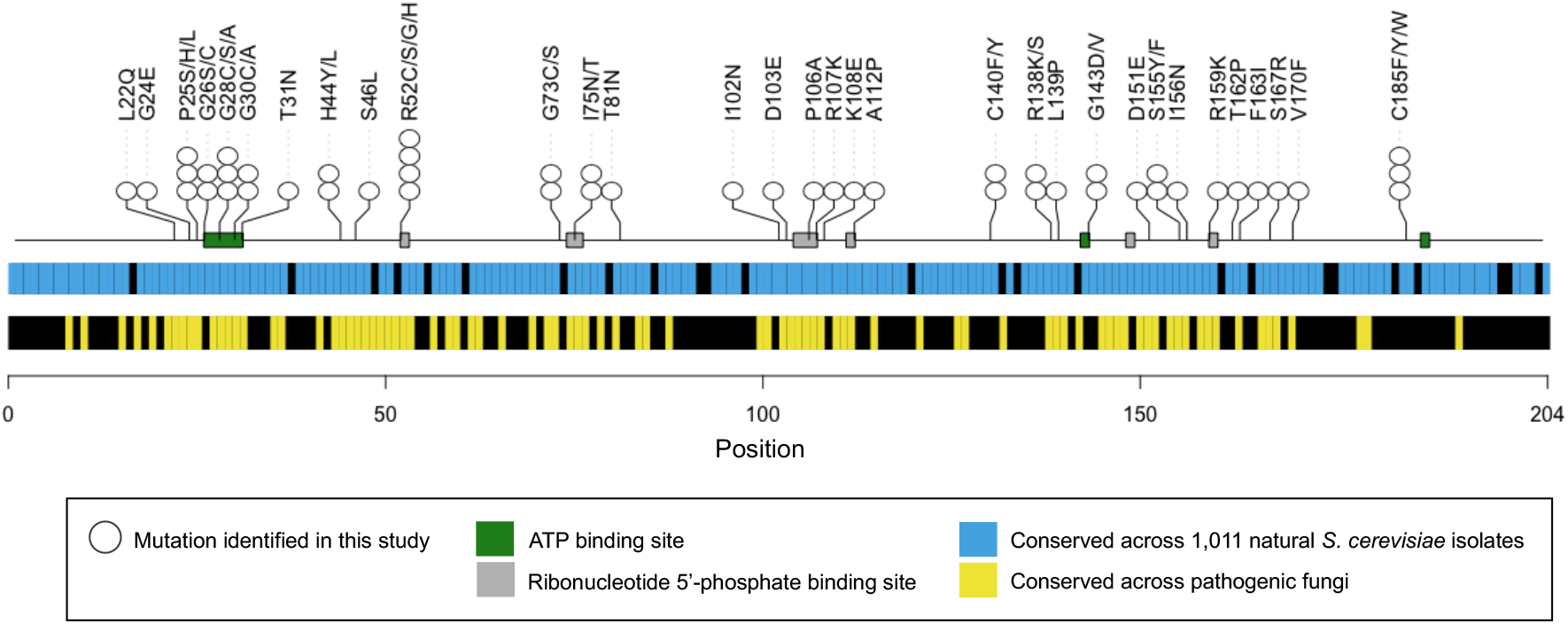
Mutations which lead to resistance to fluorinated pyrimidine derivatives occur in evolutionarily conserved regions of *URA6*. White circles denote the missense Ura6 mutations identified in this study which confer resistance to fluorinated pyrimidine derivatives. Annotated binding sites are shown in green (ATP) and in gray (ribonucleotide 5’-phosphate). Conserved Ura6 residues between 1,011 natural *S. cerevisiae* isolates are shown in blue. Conserved residues between *S. cerevisiae* (S288C), *A. fumigatus* (Af293), *C. neoformans* (JEC21), and *C. albicans* (SC5314) are shown in yellow.

We then compared the positions of *ura6* mutants in our study to the variation in *URA6* among 1,011 wild isolates of *S. cerevisiae* strains (Peter et al. 2018). All but one of the mutations we identified occur in residues that are conserved across all 1011 natural yeast isolates (Fig. 2, Supplementary Figure 9). We further compared the Ura6 amino acid sequence between *Saccharomyces cerevisiae* (S288C) and three clinically relevant pathogenic fungi, *Aspergillus fumigatus* (Af293), *Crytopcoccus neoformans* (JEC21), and *Candida albicans* (SC5314) using Ura6 protein sequences from FungiDB (Basenko et al. 2018). We found that *ura6* mutants are significantly enriched in residues that are conserved among all four species, with 28 of the 32 mutation sites in conserved residues (*p* = 1.866e-0.8, Fisher’s exact test, Fig. 2). Taken together, these mutations are located in residues that are likely to be evolutionarily and functionally important.

### UMP accumulation drives resistance in ura6 mutants

*URA6* is an essential gene, so we predicted that the 50 missense mutations we identified do not result in complete loss of function of the enzyme. We hypothesized that these mutations reduce Ura6 activity leading to an accumulation of the substrate UMP. In the presence of the fluorinated substrates, UMP and 5-FUMP both compete for the Ura6 active site. We hypothesized that elevated UMP concentrations lead UMP to be preferentially processed over 5-FUMP, reducing the conversion of 5-FUMP to 5-FUDP and limiting the production of downstream cytotoxic metabolite, 5-FUTP. To test this hypothesis, we selected two *ura6* mutants, D151E and C185W, that exhibited minimal growth reduction in the presence of fluorinated pyrimidines, and cultured them alongside WT cells in minimal media with and without 5-FOA. We measured intracellular metabolite pools in biological triplicate alongside media-only controls using LC-MS (Fig. 3A).

**Figure 3.**
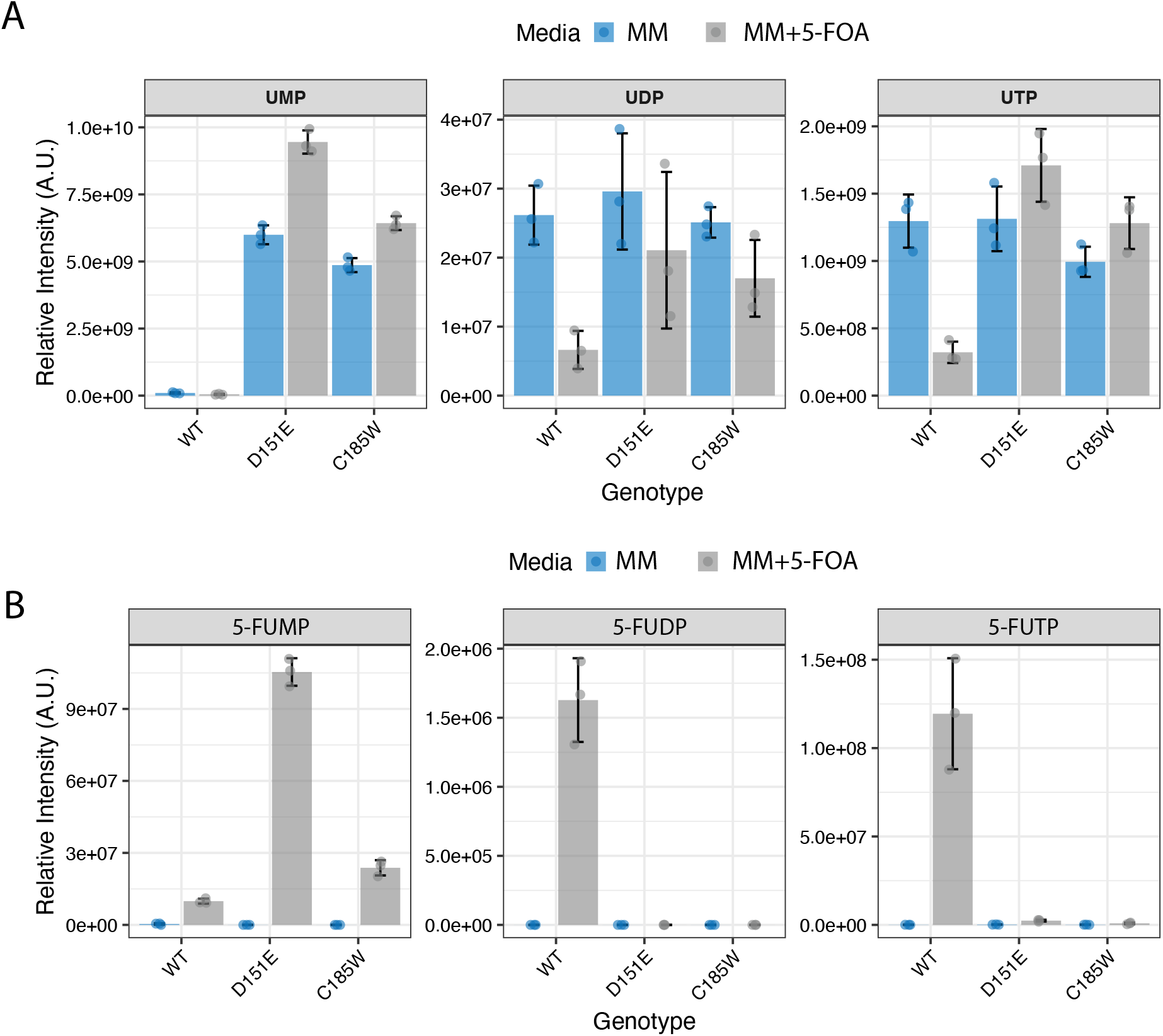
Comparative abundance of metabolites in the uracil pathway between wild-type and two *ura6* mutants (D151, C185W). A) Concentrations of UMP, UDP, and UTP and B) 5-FUMP, 5-FUDP, and 5-FUTP were measured in biological triplicate for wild type cells and two *ura6* mutants (D151E and C185W) cultivated in minimal media (blue) and in minimal media containing 5-FOA (grey) using LC-MS. Due to the differences in ionization efficiencies, it is not possible to compare intensities across analytes and relative intensities should be compared within individual analytes.

In MM, both *ura6* mutants contained higher levels of UMP than WT cells (Fig. 3A), which is consistent with a reduction in Ura6 kinase activity. While UMP levels were markedly elevated in *ura6* mutants compared to WT, the levels of the downstream metabolites, UDP and UTP, were not correspondingly reduced. UDP and UTP pools remained near wild-type levels despite a >20-fold increase in the UMP/UDP ratio (Supplementary Figure 10). This indicates that the residual Ura6 activity in the mutants is sufficient to sustain UDP/UTP homeostasis.

In MM + 5-FOA, both UMP and 5-FUMP levels were elevated in *ura6* mutants relative to WT cells (Fig. 3B). We found that WT cells accumulated substantially more 5-FUTP than *ura6* mutants, supporting the hypothesis that elevated UMP outcompetes 5-FUMP for the limited Ura6 enzyme, thereby reducing the flux of fluorinated metabolites toward the toxic downstream product, 5-FUTP. We found that uracil, uridine, and cytidine were also elevated in *ura6* mutants, consistent with a backflux through the nucleoside salvage pathway when forward conversion of UMP to UDP is impaired. Several upstream *de novo* pathway intermediates (N-carbamoyl-aspartate, dihydroorotate, orotidine, and PRPP) were elevated 8-to 30-fold in *ura6* mutants on 5-FOA, indicating that the UMP-to-UDP bottleneck also propagates back through the *de novo* pathway (Supplementary Figure 11).

We additionally observed that 5-FdUMP, a second cytotoxic metabolite derived from 5-FUMP which inhibits thymidylate synthase, was reduced ∼90 fold in both *ura6* mutants on 5-FOA, which is comparable to the 5-FUTP reduction (Supplemental Figure 9). Consistent with the loss of 5-FdUMP in *ura6* mutants, dUMP itself remains at baseline levels rather than accumulating as it does in WT cells on 5-FOA indicating that thymidylate synthase activity is preserved in the mutants. These observations indicate that the *ura6* block prevents accumulation of both known cytotoxic species of 5-FOA (5-FUTP and 5-FdUMP) simultaneously.

Combining the genetic and metabolomic data, we propose a working model for the relationship between Ura6 activity and 5-FOA sensitivity (Fig. 4). In WT cells with intact Ura6 activity, normal pyrimidine synthesis will be performed to produce UTP and CTP for basic cellular functions. In the presence of 5-FOA, Ura6 will also catalyze 5-FUMP into 5-FUDP and generating toxic 5-FUTP and 5-FCTP, largely reducing cellular viability (Fig. 4A). In *ura6* mutants in the absence of the fluorinated compounds, reduced activity of Ura6 leads to accumulation of UMP in the cell, upregulation of its conversion to uracil and further export (Fig. 4B). Based on the reduced Ura6 activity, we would predict a lower level of downstream metabolites of UDP and UTP. However, similar levels of UDP and UTP were observed in *ura6* mutants compared with WT. This suggests that there may be cellular homeostasis mechanisms that compensate the UDP and UTP to the WT level in *ura6* mutants. In the presence of 5-FOA, the reduced activity of Ura6 will also reduce the production of 5-FUDP and accumulation of 5-FUMP. The fact that we do not observe quantifiable level of 5-FUDP in *ura6* mutants suggests that UMP may outcompete 5-FUMP for the Ura6 enzymatic function, inhibiting the production of 5-FUDP and therefore downstream metabolites.

**Figure 4.**
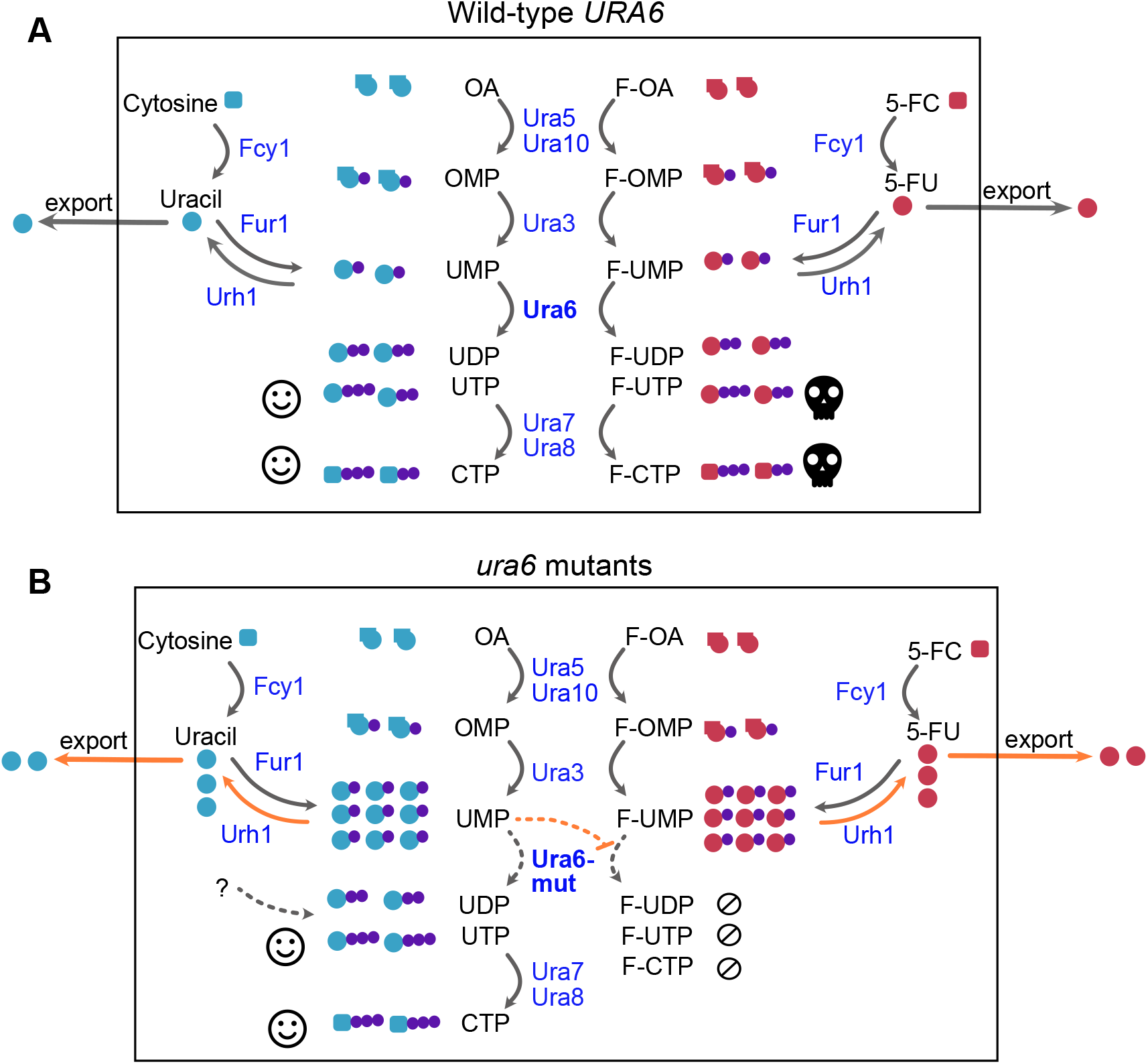
Working model for fluorinated pyrimidine resistance of *ura6* mutants. **A**) Wild type cells metabolize both OA and 5-FOA. The conversion of 5-FOA into 5-FUTP and 5-FCTP is toxic and leads to cell death. B) *ura6* mutants are resistant to 5-FOA (and other fluorinated pyrimidine derivatives) because they do not convert 5-FOA into 5-FUTP and 5-FCTP. Instead, *ura6* mutants accumulate both UMP and 5-FUMP. UMP is preferentially metabolized by Ura6 to generate enough UTP and CTP for the cell to grow, and very little 5-FUMP is metabolized into the toxic downstream products 5-FUTP and 5-FCTP. The accumulation of UMP and 5-FUMP results in the conversion to uracil and 5-FU, which is exported from the cell.

## Discussion

Mutations to *URA6* can lead to a reduction of uridylate kinase activity and confer resistance to fluorinated pyrimidine derivatives. This finding suggests that previously uncharacterized *URA3+* mutants observed to have 5-FOA resistance could be due to mutations in *URA6*. One reason this may have evaded previous studies could be due to the experimental design. Most 5-FOA-based selection for *URA3* is on higher frequency events, such as plasmid loss or mitotic recombination, on the order of 10^-5^ events per genome per cell division or higher (e.g., Jinks-Robertson and Petes 2021), which won’t be affected by the low frequency of the *ura6* point mutants. Our study plated a large number of cells – on the order of 10^8^ for each sample, including the overnight inoculant. By contrast, standard fluctuation-based mutation assays dilute 10,000 fold from the overnight inoculant, making it less likely to observe any mutants that arose during the overnight incubation. In addition, we used media depleted of uracil to grow overnight cultures, which selects against *ura3* mutants but not *ura6* mutants. Future studies should consider mutations to *URA6* when plating large quantities of cells to select for *URA3* using 5-FOA, especially when cells are pre-grown in uracil dropout media. In contrast, *CAN1* appears to account for 100% of canavanine-resistant mutants (Lang and Murray 2008; Jiang et al. 2021). As an example, among 227 mutants that confer canavanine resistance, all of them have a mutation in the *CAN1* after individual Sanger sequencing (Lang and Murray 2008). 5-FOA resistance is distinct from canavanine resistance in that there is a secondary mutable target, *URA6*, which unlike *URA3*, cannot be excluded by uracil-dropout selection.

In this study, we identified 50 unique missense mutations to Ura6 that confer resistance to fluorinated pyrimidines. The *ura6* mutants we identified tend to reside in conserved regions across pathogenic yeasts across large evolutionary distances. Quantitatively, we observed that WT cells grow slightly better in MM compared to *ura6* cells, which means *ura6* cells will be outcompeted against in non-fluorinated conditions. Most of the mutations we found occur in sites that are conserved across *S. cerevisiae* and pathogenic fungi, however we also found four mutation sites which confer resistance in *S. cerevisiae* are located in non-conserved regions. We found that *ura6* mutants do not show an obvious growth trade-off in the presence or absence of fluorinated compounds, as previously seen in a study of mutations to *FCY1 (Després et al. 2022)*. The fact that *ura6* mutants can grow in MM, 5-FC, 5-FU, and 5-FOA, places them in a unique category in comparison to WT or *ura3* cells, which can only grow in a subset of these conditions (Fig. 1A). Mutants like *ura6* may be an example of a stepping stone for adaptation, where a compromise is not a requirement when growing in a novel environment (Wang et al. 2024); however, it is also possible that we have not explored all plausible environments in which *ura6* can show the trade-off.

The accumulation of UMP we observed in our *ura6* mutants is consistent with reduced uridylate kinase activity, which could be the consequence of reduced levels or activity of the mutated Ura6 protein. The broad distribution of 5-FOA resistant *ura6* mutations across the protein sequence is inconsistent with a mechanism specific to substrate binding, since many mutations are located in domains not associated with binding to the UMP substrate. Instead, we suggest that the decreased production of FUTP in these *ura6* mutants is most likely an example of higher affinity binding of UMP over FUMP. In studies of mammalian and bacterial uridylate kinases, UMP is a higher affinity substrate than FUMP (Humeniuk et al., 2009, Gagyi et al., 2003). Intracellular concentrations of UMP in *S. cerevisiae* grown on similar conditions have been observed as high as 145 µM (Park et al., 2016), in range of the K_m_ observed across uridylate kinases of other organisms. Assuming complete conversion of FOA to FUMP, the intracellular concentration of FUMP in our experiments would be approximately 8 mM, in substantial excess of the physiological range of wild type cells. However, in the context of the *ura6* mutations that increase UMP levels by as much as 140 to 206-fold in 5-FOA media (Fig. 3A), concentrations of UMP would likely exceed those of FUMP. In this scenario, the affinity preference of the Ura6 enzyme for UMP over FUMP, in combination with the continual turnover of UDP and UTP, would result in nearly exclusive occupancy of Ura6 on UMP as compared to FUMP, consistent with our metabolomic observations of extremely low levels of FUDP or FUTP in *ura6* mutants.

Recently, a study by Kowal and colleagues reported a similar observation, in which *ura6* mutants were able to grow on 5-FOA in the absence of uracil in fission yeast *Schizosaccharomyces pombe* (Kowal et al. 2026). In that work, the mutants were first identified in mutation assays using quiescent cells under 5-FOA selection and were later confirmed in actively growing cells. Our genetic and growth experiments complement their findings and our metabolomic data further characterize the mechanism connecting mutations to *URA6* to 5-FOA resistance. Uridylate kinase activity is essential and conserved across fungi and mammals (the human homolog is CMPK1, Alliance of Genome Resources Consortium 2020). The reduction of uridylate kinase activity appears to be a conserved route to fluoropyrimidine resistance across species.

## Data Availability Statement

Strains are available upon request. Supplementary table 1 contains a list of all mutations identified in this work. Raw sequence data is available on the SRA BioProject: PRJNA1514863. Data for growth rate and metabolic analysis are available in the supplemental material.

## Acknowledgment

JOA acknowledges support from University of Washington Biological Mechanisms of Healthy Aging Training Grant T32AG066574. PJ was supported by a Burroughs Wellcome Fund Career Award at the Scientific Interface awarded to KH. KH acknowledges additional support from a Searle Scholarship, a Sloan Research Fellowship, a Pew Biomedical Scholarship, and National Institute of General Medical Sciences Grant 1R35GM133428-01. MD acknowledges support from the Nathan Shock Centers of Excellence in the Basic Biology of Aging, P30AG013280. The content is solely the responsibility of the authors and does not necessarily represent the official views of the Burroughs Wellcome Fund, the Kinship Foundation, the Sloan Foundation, the Pew Charitable Trust, the NIH, or NIGMS. We also acknowledge the help of Ryan Sheldon at the Van Andel Institute Mass Spectroscopy core (RRID:SCR_024903) for his work on measuring metabolites using LC-MS. MJD holds the William H. Gates III Endowed Chair in Biomedical Sciences.

## Materials and Methods

### Strains

FY4 *MAT*α*flo1::KanMx, flo9::NatMx*

#### Media

Minimal media (MM): Yeast nitrogen base, without amino acids and without ammonium sulfate (1.7g/L), Ammonium Sulfate (5g/L), Dextrose (20g/L)

Minimal media + fluorinated prodrug (MM+5-FC, MM+5-FU, MM+5-FOA): A concentration of 1mg/mL was used for all fluorinated prodrug media.

Synthetic complete media (SC): Yeast nitrogen base, without amino acid and without ammonium sulfate (1.7g/L), Ammonium Sulfate (5g/L), Dextrose (20g/L), Adenine (0.019g/L), Arginine (0.019g/L), Aspartic Acid (0.096g/L), Glutamic Acid (0.096g/L), Histidine (0.019g/L), Isoleucine (0.077g/L), Leucine (0.077g/L), Lysine (0.058g/L),

Methionine (0.019g/L), Phenylalanine (0.048g/L), Serine (0.384g/L), Threonine (0.192g/L), Tryptophan (0.077g/L), Tyrosine (0.058g/L), Uracil (0.019g/L), Valine (0.144g/L)

Synthetic complete media without uracil (SC-ura): Yeast nitrogen base, without amino acid and without ammonium sulfate (1.7g/L), Ammonium Sulfate (5g/L), Dextrose (20g/L), Adenine (0.019g/L), Arginine (0.019g/L), Aspartic Acid (0.096g/L), Glutamic Acid (0.096g/L), Histidine (0.019g/L), Isoleucine (0.077g/L), Leucine (0.077g/L), Lysine (0.058g/L), Methionine (0.019g/L), Phenylalanine (0.048g/L), Serine (0.384g/L), Threonine (0.192g/L), Tryptophan (0.077g/L), Tyrosine (0.058g/L), Valine (0.144g/L)

### ura6 *mutant discovery and sequencing analysis*

The original experiment was set up to use *URA3* to quantify mutation rates across the lifespan of yeast. Overnight inoculants were grown in SC-ura media, which were subsequently labeled with magnetic beads. Continuous cultures with magnets are set up to capture the mother cells, under three conditions, SC-ura without glucose, SC, and SC-ura. Mother and daughter cells from different time points (day 0, day 1, day 4, day 6) were harvested and tested for their mutation frequencies using 5-FOA resistance selection.

Generating the mutant pools follows the procedure in (Jiang et al. 2021; Jiang, Ollodart, and Dunham 2022), pooling roughly 35 mutants per pool. We used PCR to amplify the *URA3* locus (*URA3* primers from Lang and Murray 2008) and prepared the PCR product for Illumina sequencing using the Nextera XT kit. We used the same mutation calling pipeline as (Jiang et al. 2021), without excluding multi-nucleotide mutations. To identify mutations to *URA6*, we used PCR to amplify the *URA6* locus (forward primer: CCACGCGCTAACAAAAGTCA, reverse primer: CAGTGCTCCCGCCATTGTTA) and prepared the PCR product for Illumina sequencing from the same pooled genomic DNA. The same mutation calling pipeline as *URA3* was used.

10 individual mutants were subjected to whole-genome sequencing using the Illumina DNA Flex Library Prep Kit and sequenced on Nextseq 550. We performed the variant calling pipeline as previously described (Taylor et al. 2022).

### Quantification of cell growth

To quantify the growth of *ura6* mutants, three individual clones were isolated from eight unique mutants grown in 200µL minimal media in 96-well plates for 24 hours at 30°C. The 24-hour cultures were resuspended and 2µL of each culture was transferred into 200µL of the following media types: synthetic complete media (SC media), synthetic media minus uracil (SC-ura), minimal media (MM), minimal media with 5-FOA (MM+5-FOA, 1mg/mL), minimal media with 5-FU (MM+5FU, 1mg/mL), and minimal media with 5-FC (MM+5FC, 1mg/mL). A Biotek Synergy H1 plate reader was used to monitor optical density every 15 minutes over the course of 48 hours. Growth measurements were conducted at both 30°C and 37°C to assess temperature sensitivity. Saturation density was taken as the median of the final 15 readings of each well. Maximum specific growth rate (μmax) was determined from the slope of ln(OD600) against time. μmax was taken as the median slope of the five steepest retained slopes.

### Conservation

To identify the natural variation of Ura6 within *Saccharomyces cerevisiae* strains, amino acid sequences of Ura6 from the 1,011 *S. cerevisiae* natural isolate collection were used to perform a within-species alignment. We used MAFFT v 7.487 (Katoh and Standley 2013) to perform multiple sequence alignments with an accurate alignment option (mafft-linsi) with default parameters. To identify conserved regions between pathogenic fungi and *Saccharomyces cerevisiae* we compared the amino acid sequence of Ura6 in *S. cerevisiae* (S288C), *A. fumigatus* (Af293), *C. neoformans* (JEC21), and *C. albicans* (SC5314) using Clustal alignments collected from from FungiDB. We only scored residues that were conserved between all four species. We used the bioconductor package, LollipopPlot (Ou and Zhu 2019) to visualize the positions of the mutations and their relative frequency and a custom R script to generate the conservation tracks.

### Metabolite extraction

Metabolite extraction was carried out as previously described by Rosebrock and Caudy (2017). To examine the metabolite pools of wild-type and *ura6* mutant cells, cells were grown in minimal media, supplemented with 0.5g/L ammonium sulfate and 2.62 g/L ammonium acetate as a nitrogen source overnight. The following day, the overnight cultures were diluted so that OD_600_ = 0.1 in either minimal media or minimal media + 5-FOA. These cells were allowed to grow for approximately 3 hours until their OD_600_ > 0.4. 5mL of culture was transferred to a filter apparatus equipped with a 0.2 micron nylon filter. Filtered cells were immediately transferred to 1.2mL Methanol:Acetonitrile:Water (2:2:1) solution and rapidly vortexed and placed in dry ice until frozen. Media only controls were included and processed identically to samples containing cells. The filter apparatus was rinsed with 15mL of water between each sample. Once all samples were processed and frozen they were moved back and forth between dry ice and a -20°C freezer for a total of 3 freeze thaw cycles. Upon the final cycle, filter papers were removed and samples were centrifuged for 10 minutes at 4°C to pellet cells. Finally, 1mL of supernatant was removed for LC-MS analysis.

### LC-MS analysis

LC-MS analysis was carried out as previously described by House and colleagues (House et al., 2024) Briefly, samples were re-extracted using a chloroform/methanol/water solvent system to maximize recovery of nucleotides and other polar metabolites. Metabolomics data were acquired by ion-pairing liquid chromatography coupled to a Thermo Scientific Orbitrap Exploris 240 mass spectrometer. Pooled unlabeled samples served as quality-control injections and were used to acquire data-dependent MS2 (ddMS2) spectra for compound identification. Targeted analysis was performed in Skyline. Metabolites were annotated on the basis of retention-time matching to authentic chemical standards and MS2 spectral library matches associated with each chromatographic method.

