## Supplementary Figures for "*URA6* mutations provide an alternative mechanism for 5-FOA resistance in *Saccharomyces cerevisiae*"

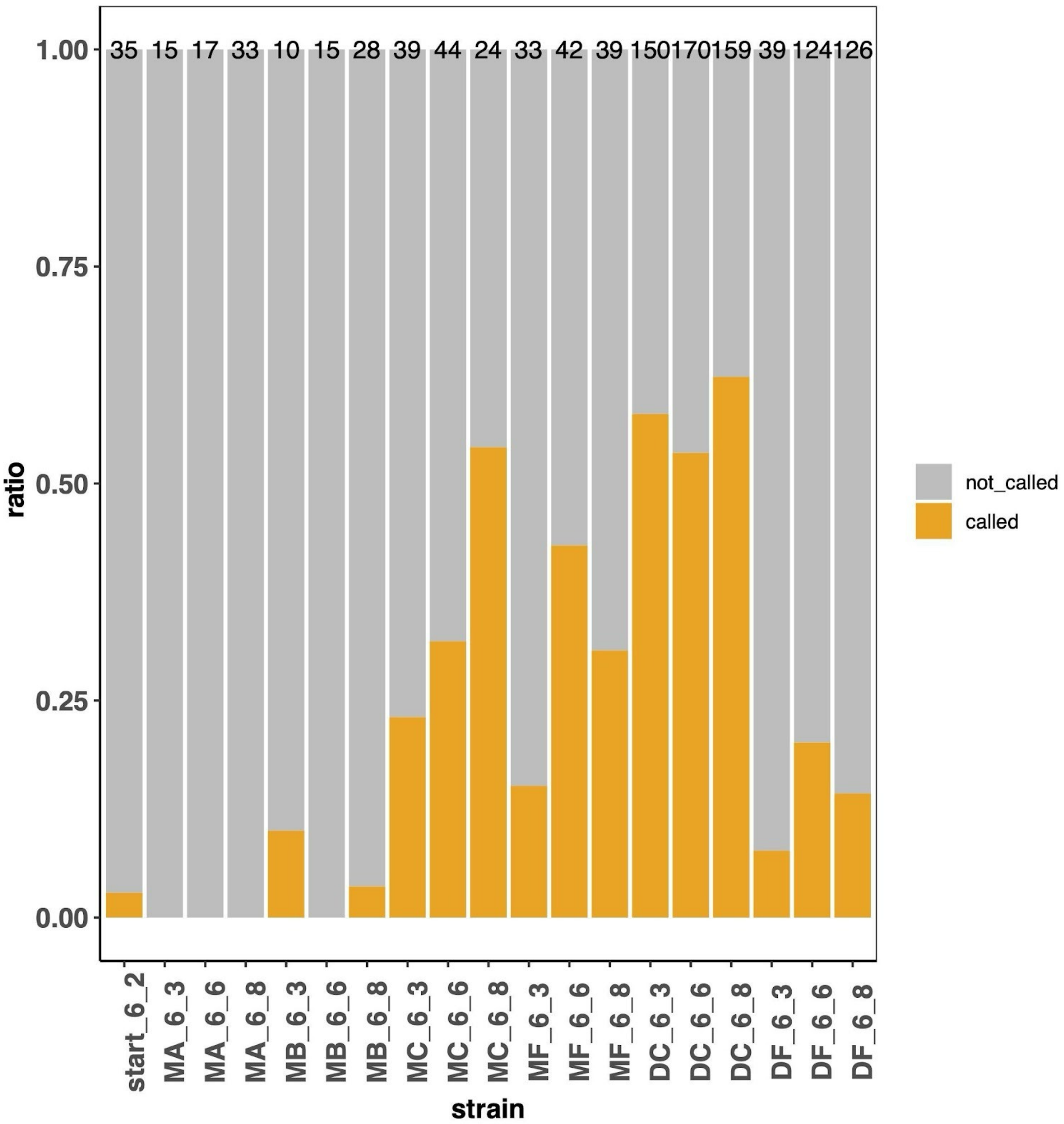


**Supplementary Figure 1.** The ratio of *ura3* mutants in each 5-FOA resistant sample. The number on the top of each bar shows the total number of individual mutants pooled in each sample. The orange bar shows the ratio of called *ura3* mutants from the computational pipeline after sequencing by the total number of mutants pooled. The two numbers in each name denote the date of the sample, for example, "6_2" means June 2. "Start" sample is the overnight inoculant. Samples beginning with "M" are mother cells, "D" are daughter cells. The second letter denotes different growth conditions, "A" and "B" are SC-ura without no glucose, "C" and "D" are SC, "E" and "F" are SC-ura.

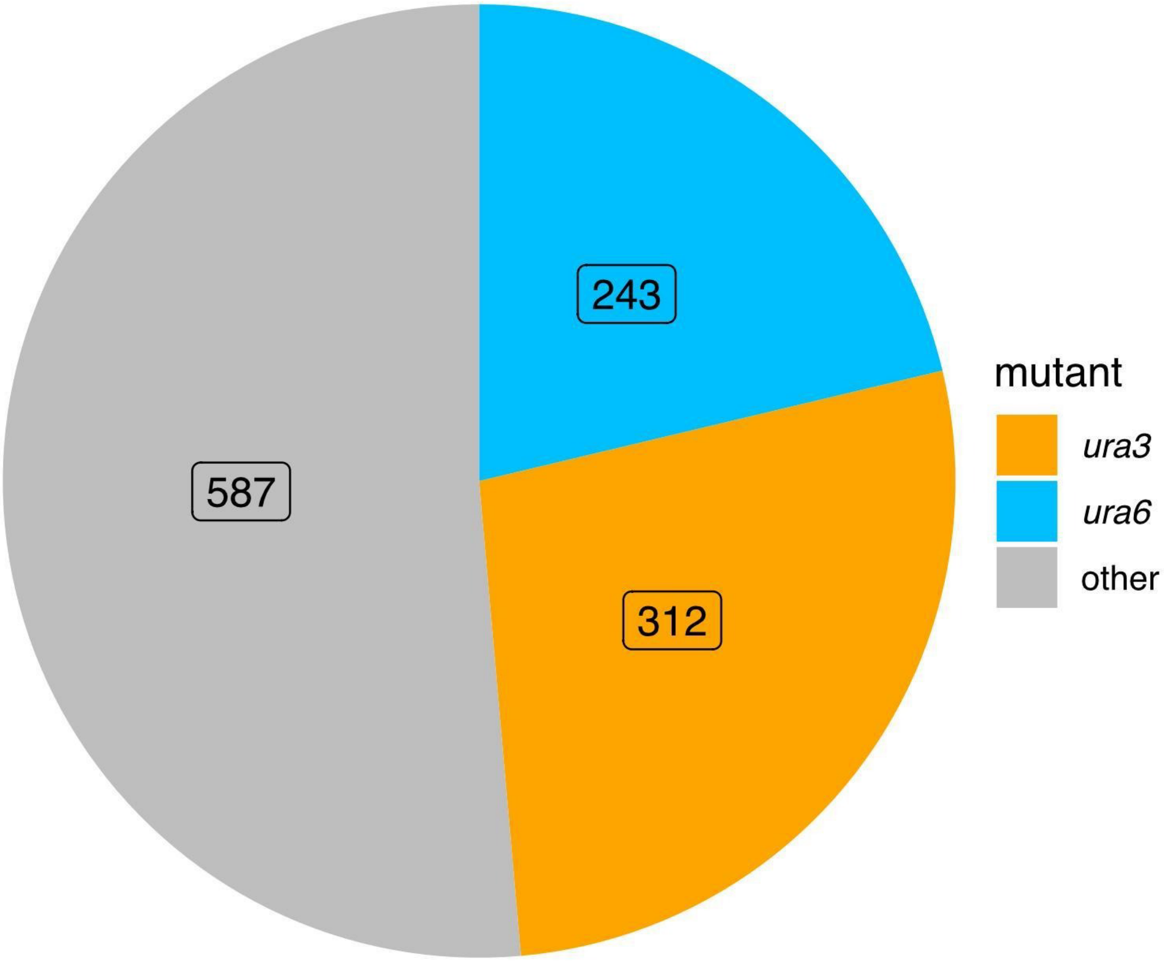


**Supplementary Figure 2.** A pie chart displaying *ura3 and ura6* mutants called across all mutant samples. Blue indicates the *ura6* mutants, and yellow indicates *ura3*. Grey indicates the remaining mutants that are pooled but not called either *ura3* or *ura6* in our pipeline.

| Sample | Position | Reference | Mutation |
|---|---|---|---|
| MC1_6_6_m10_S21 | 75 | I | T |
| MC1_6_6_m1_S19 | 75 | I | T |
| MC1_6_6_m18_S22 | 155 | S | F |
| MC1_6_6_m3_S20 | 155 | S | F |
| MC1_6_6_m19_S23 | 31 | T | N |
| MB_6_3_m2_S14 | 31 | T | N |
| MB_6_3_m6_S16 | 31 | T | N |
| MB_6_3_m7_S17 | 31 | T | N |
| MB_6_3_m8_S18 | 31 | T | N |
| MB_6_3_m5_S15 | 143 | G | D |

**Supplemental Table 1.** Missense mutations identified in Ura6 via whole genome sequencing

| Position | Reference | Mutation |
|---|---|---|
| 22 | L | Q |
| 24 | G | E |
| 25 | P | S |
| 25 | P | H |
| 25 | P | L |
| 26 | G | S |
| 26 | G | C |
| 28 | G | C |
| 28 | G | S |
| 28 | G | A |
| 30 | G | C |
| 30 | G | A |
| 31 | T | N |
| 44 | H | Y |
| 44 | H | L |
| 46 | S | L |
| 52 | R | C |
| 52 | R | S |
| 52 | R | G |
| 52 | R | H |
| 73 | G | C |
| 73 | G | S |
| 75 | I | N |
| 75 | I | T |
| 81 | T | N |
| 102 | I | N |
| 103 | D | E |
| 106 | P | A |
| 107 | R | K |
| 108 | K | E |
| 112 | A | P |
| 130 | C | F |
| 130 | C | Y |
| 138 | R | K |
| 138 | R | S |
| 139 | L | P |
| 143 | G | D |
| 143 | G | V |
| 151 | D | E |
| 155 | S | Y |
| 155 | S | F |
| 156 | I | N |
| 159 | R | K |
| 162 | T | P |
| 163 | F | I |
| 167 | S | R |
| 170 | V | F |
| 185 | C | F |
| 185 | C | Y |
| 185 | C | W |

**Supplementary Table 2.** Complete list of the 5-FOA resistant, missense mutations to Ura6 identified in this study.

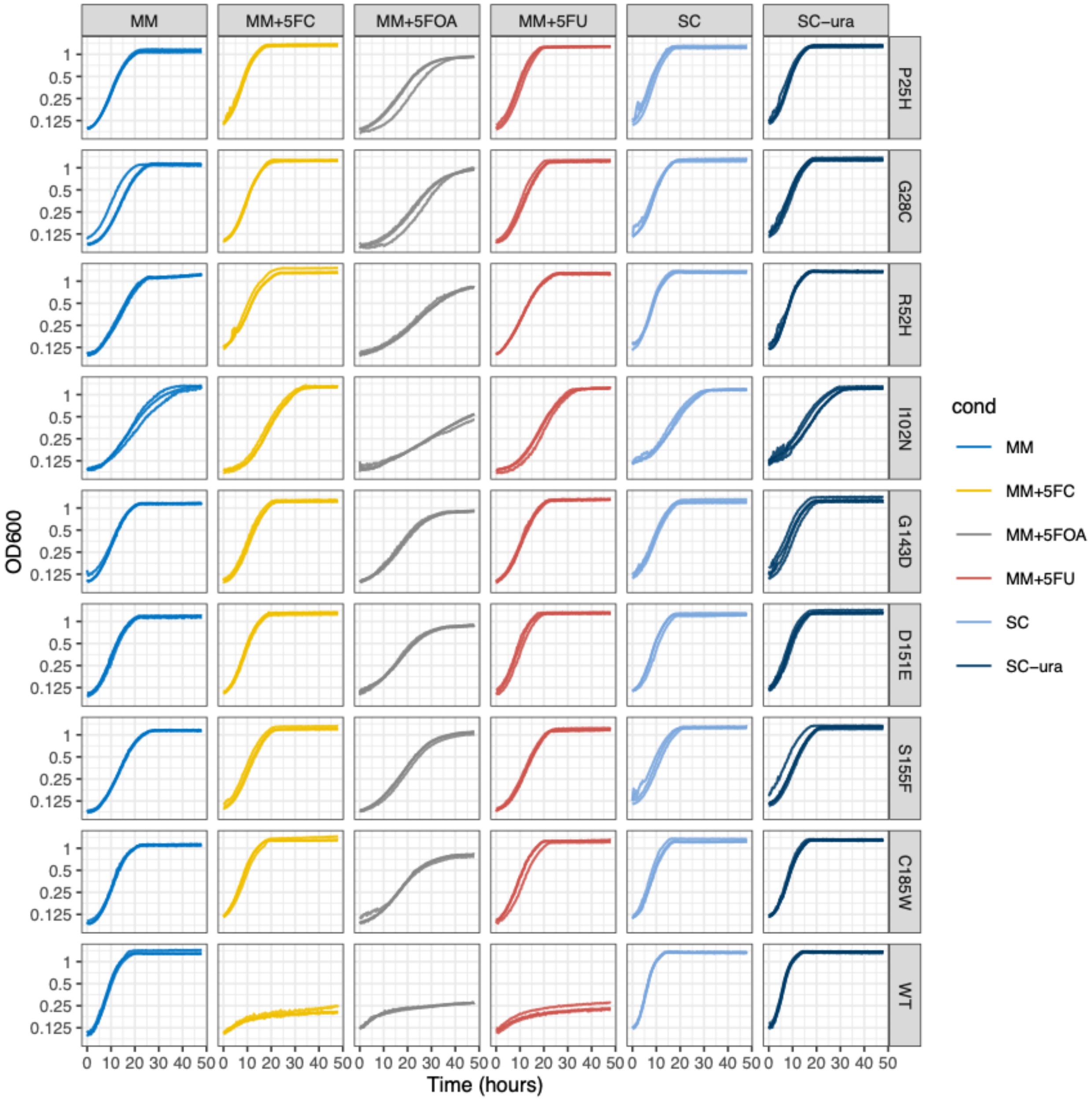


**Supplementary Figure 3.** Growth curves of wild type and *ura6* cultures grown in MM, MM+5FC, MM+5-FOA, MM+5FU, SC, and SC-ura media at 30°C.

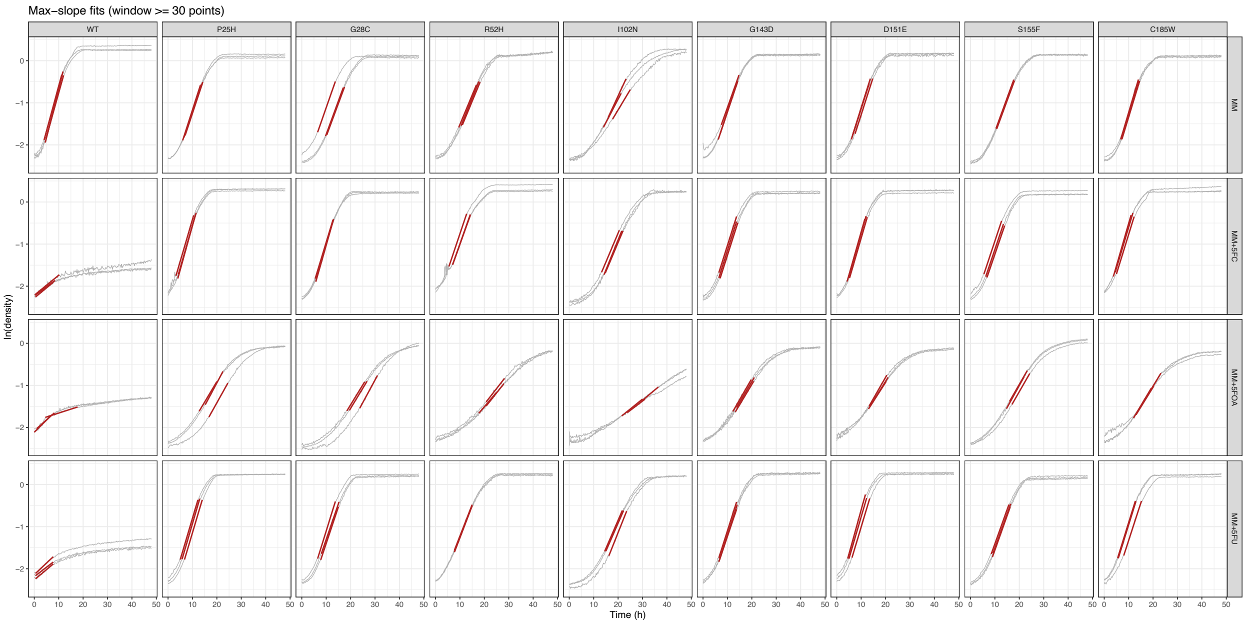


**Supplementary Figure 4.** Region of growth curve used for maximum growth rate ($\mu_{\text{max}}$) calculation.

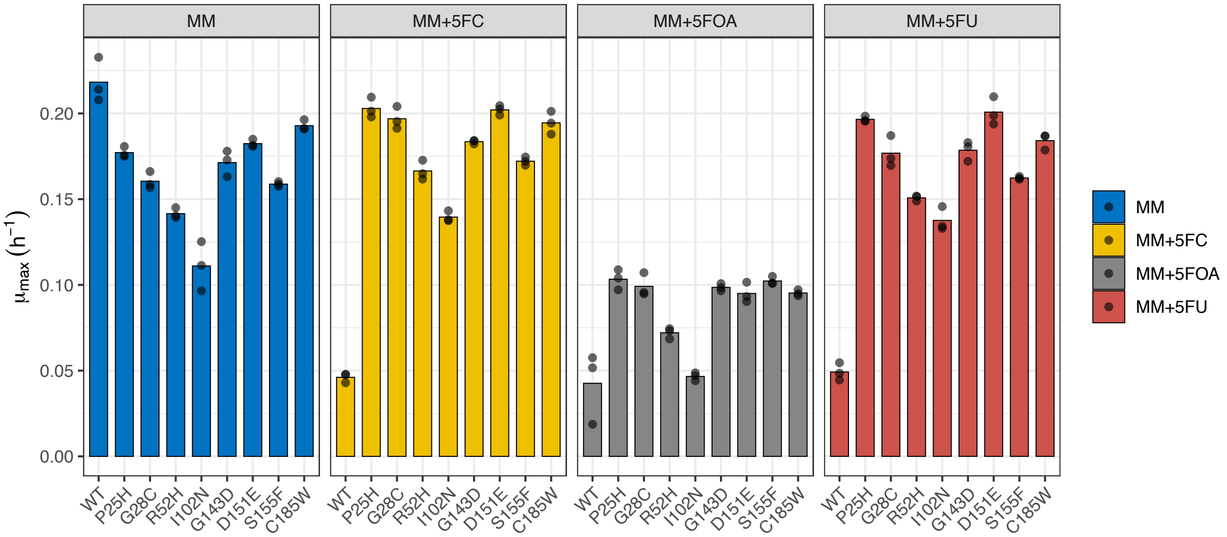


**Supplementary Figure 5.** Maximum growth rate ($\mu_{max}$) measurements of WT and *ura6* mutants in minimal media (MM), minimal media + 5-FC (MM+5FC), minimal media + 5-FOA (MM+5FOA), and minimal media + 5-FU (MM+5FU).

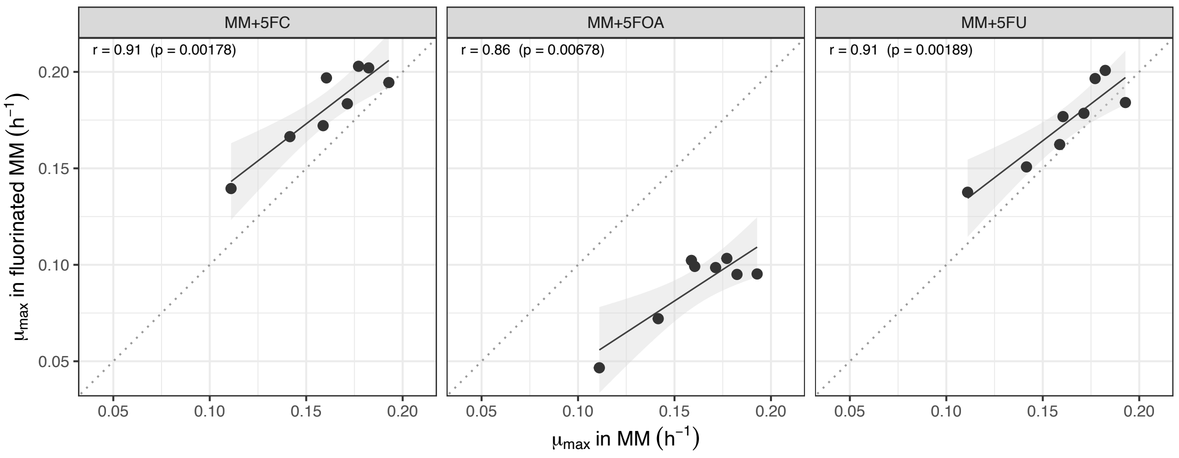


**Supplementary Figure 6.** Linear correlations of maximum growth rate ($\mu_{\max}$) measurements between MM and fluorinated drug containing media for *ura6* mutants.

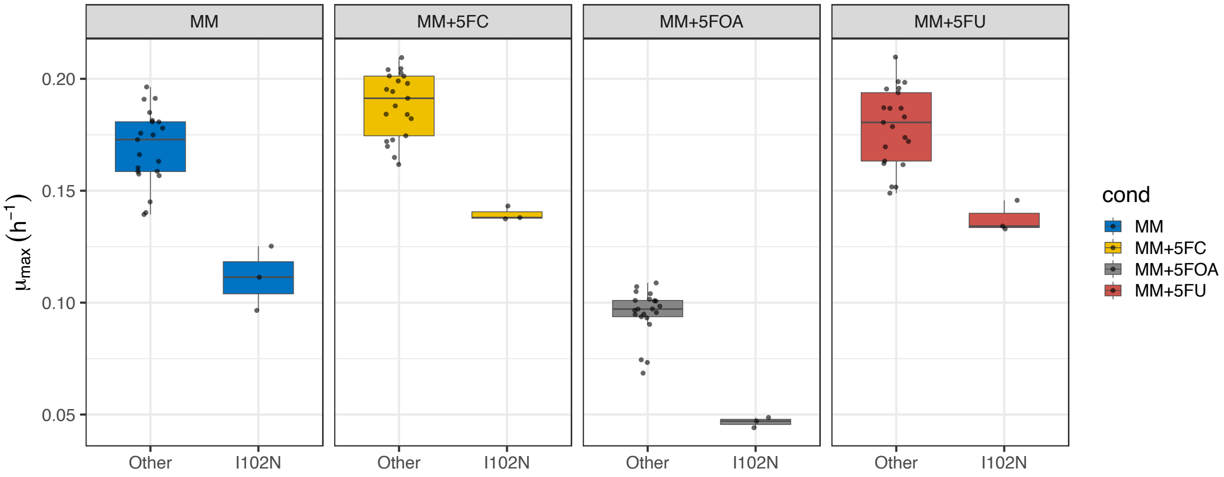


**Supplementary Figure 7 :** The temperature sensitive I102N mutant grows less robustly than the other isolated mutants in all media types. The difference in maximum growth between the I102N mutant and the rest of the *ura6* mutants was determined to be statistically significant across all media types (Welch's two sample T test, MM (p-value < 0.01 for all comparisons.)

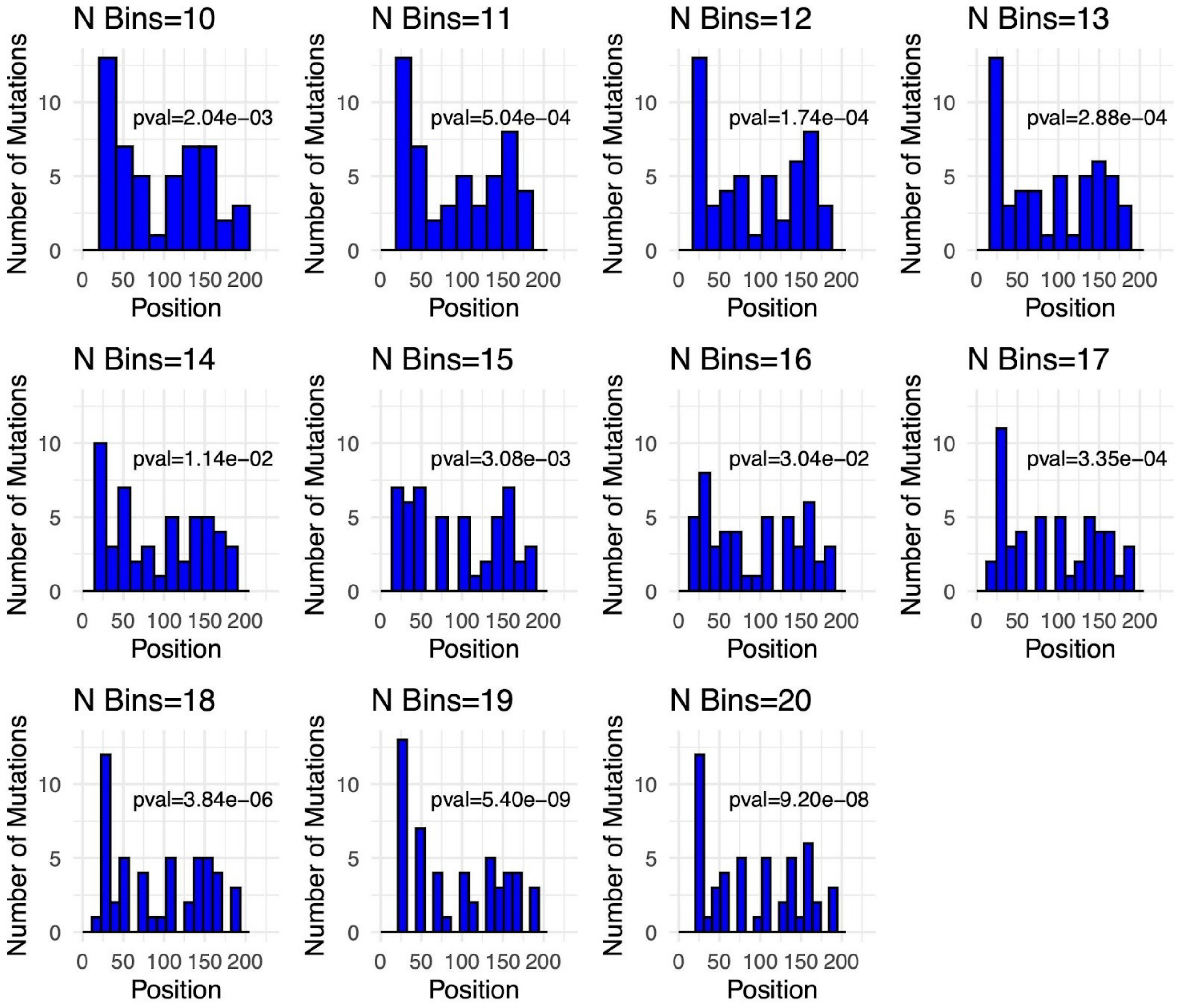


**Supplementary Figure 8.** Testing the uniformity of *ura6* missense mutation distribution along the gene. Different bin sizes are tested. Ura6 sequence is divided into different bins. Each plot shows the number of unique mutants that are present in each bin, given the bin size. The chi-square tests are performed on the observed number of mutants per bin against the expected number of mutants if they are uniformly distributed. The *p-value* is shown for each bin size. An enrichment near position 25 is observed in most of the cases.

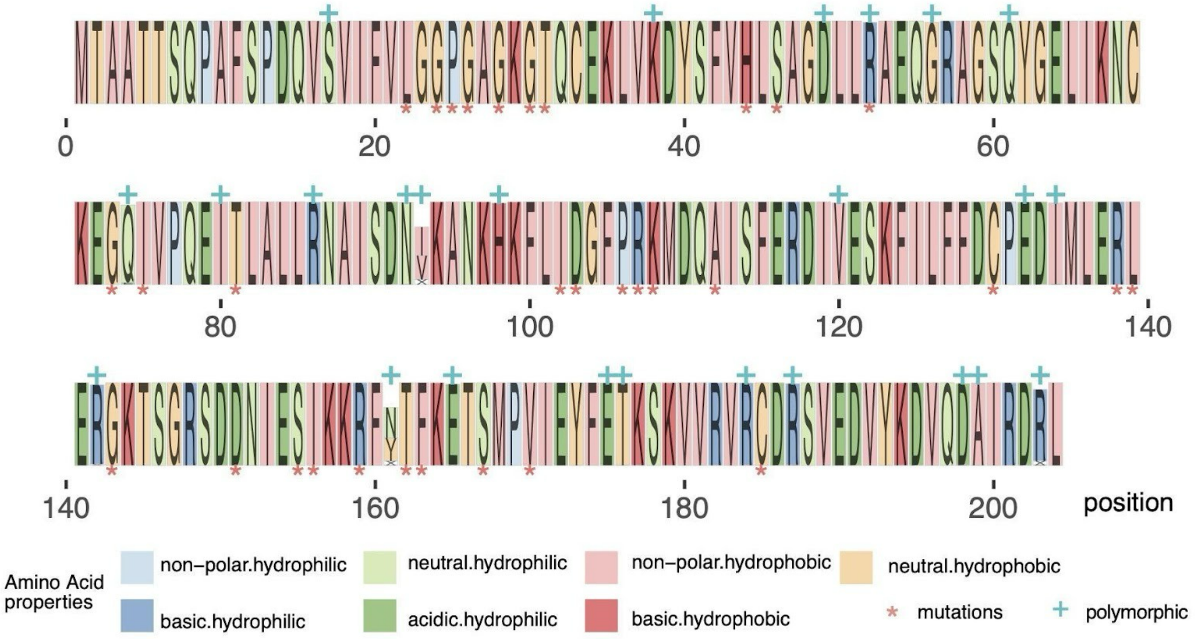


**Supplementary Figure 9.** The positions of unique missense mutations to *URA6* identified in this study compared with polymorphic sites in the 1011 *S. cerevisiae* collection. Logo plot (gglogo package(Hofmann, n.d.)) for the entire Ura6 based on all the polymorphisms in the 1011 *S. cerevisiae*. Sites that are polymorphic in the 1011 *S. cerevisiae* are marked with blue crosses. Unique *ura6* sites in our dataset are marked with red asterisks.

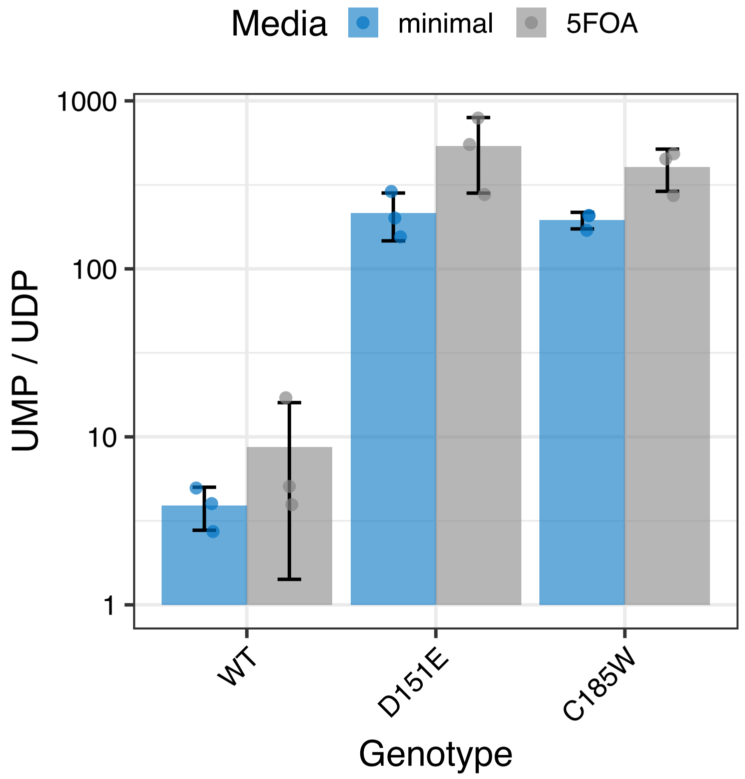


**Supplementary Figure 10.** UMP/UDP ratios for Wild Type (WT) and two *ura6* mutants (D151E and C185W). Ratios were calculated based off of the relative intensity of each metabolite for three replicates.

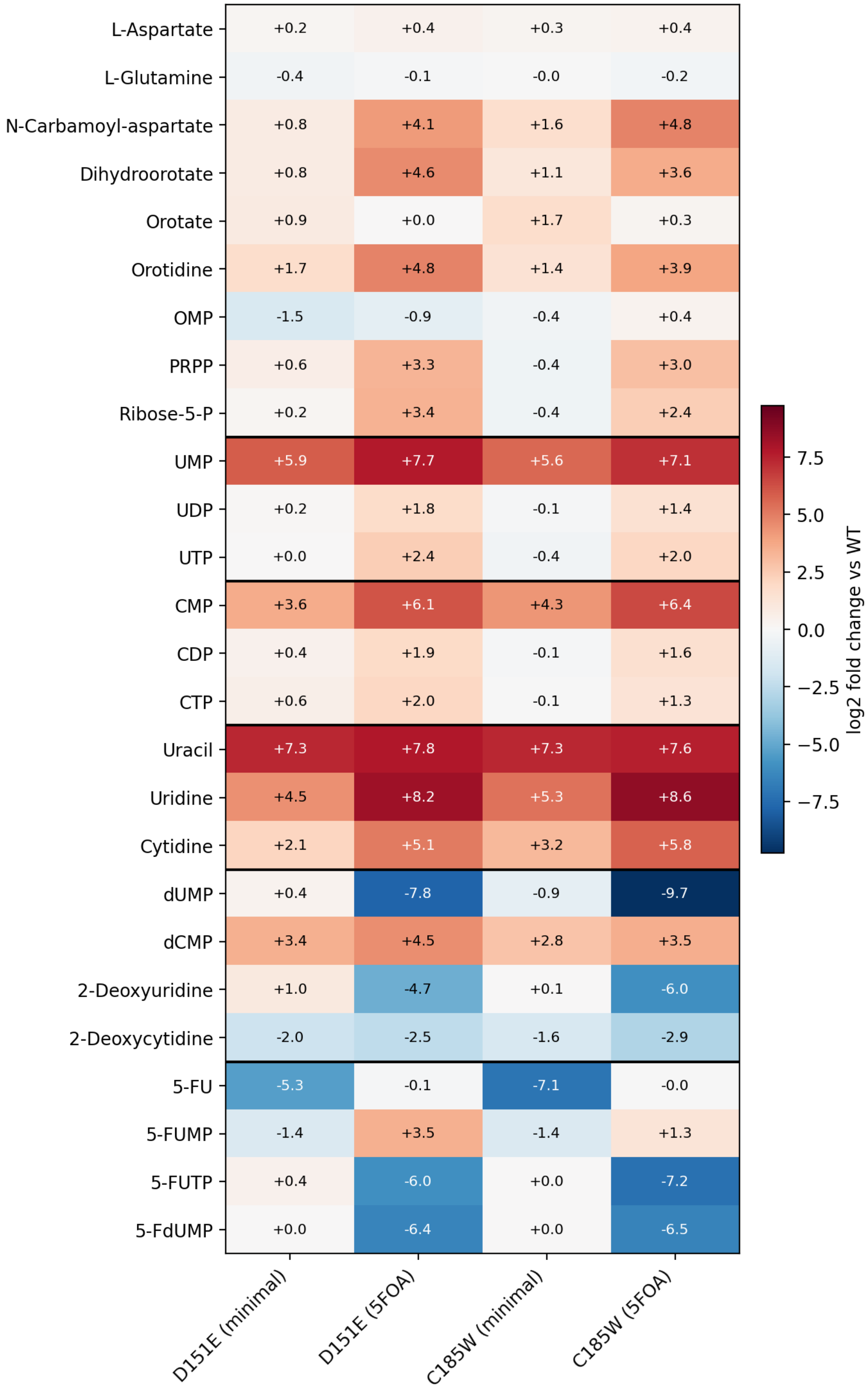

L-Aspartate
+0.2
+0.4
+0.3
+0.4
L-Glutamine
-0.4
-0.1
-0.0
-0.2
N-Carbamoyl-aspartate
+0.8
+4.1
+1.6
+4.8
Dihydroorotate
+0.8
+4.6
+1.1
+3.6
Orotate
+0.9
+0.0
+1.7
+0.3
Orotidine
+1.7
+4.8
+1.4
+3.9
OMP
-1.5
-0.9
-0.4
+0.4
PRPP
+0.6
+3.3
-0.4
+3.0
Ribose-5-P
+0.2
+3.4
-0.4
+2.4
UMP
+5.9
+7.7
+5.6
+7.1
UDP
+0.2
+1.8
-0.1
+1.4
UTP
+0.0
+2.4
-0.4
+2.0
CMP
+3.6
+6.1
+4.3
+6.4
CDP
+0.4
+1.9
-0.1
+1.6
CTP
+0.6
+2.0
-0.1
+1.3
Uracil
+7.3
+7.8
+7.3
+7.6
Uridine
+4.5
+8.2
+5.3
+8.6
Cytidine
+2.1
+5.1
+3.2
+5.8
dUMP
+0.4
-7.8
-0.9
-9.7
dCMP
+3.4
+4.5
+2.8
+3.5
2-Deoxyuridine
+1.0
-4.7
+0.1
-6.0
2-Deoxycytidine
-2.0
-2.5
-1.6
-2.9
5-FU
-5.3
-0.1
-7.1
-0.0
5-FUMP
-1.4
+3.5
-1.4
+1.3
5-FUTP
+0.4
-6.0
+0.0
-7.2
5-FdUMP
+0.0
-6.4
+0.0
-6.5
D151E (minimal)
D151E (5FOA)
C185W (minimal)
C185W (5FOA)
7.5
5.0
2.5
0.0
−2.5
−5.0
−7.5
log2 fold change vs WT

**Supplementary Figure 11. Pyrimidine pathway metabolite log2 fold-change heatmap.** Heatmap of log2-transformed fold-changes in metabolite relative intensity between *ura6* mutants and wild type, derived from untargeted LC-MS metabolomics of cells grown in minimal media (MM) or minimal media supplemented with 5-fluoroorotic acid (MM + 5-FOA). Fold-changes were calculated as the ratio of mean mutant intensity to mean wild-type intensity ($n = 3$ biological replicates per condition) within each media. Red indicates metabolites elevated in the mutant relative to wild type (positive log2 fold-change), blue indicates metabolites depleted in the mutant relative to wild type (negative log2 fold-change), and white indicates no change (log2 fold-change $\approx 0$). Numerical log2 fold-change values are overlaid on each cell. Abbreviations: UMP/UDP/UTP, uridine mono-/di-/triphosphate; CMP/CDP/CTP, cytidine mono-/di-/triphosphate; dUMP/dCMP, 2'-deoxyuridine/cytidine monophosphate; OMP, orotidine 5'-monophosphate; PRPP, 5-phosphoribosyl-1-pyrophosphate; 5-FU, 5-fluorouracil; 5-FUMP/5-FUTP, 5-fluorouridine mono-/triphosphate; 5-FdUMP, 5-fluoro-2'-deoxyuridine monophosphate.
